# Utilizing trait networks and structural equation models as tools to interpret multi-trait genome-wide association studies

**DOI:** 10.1101/553008

**Authors:** Mehdi Momen, Malachy T. Campbell, Harkamal Walia, Gota Morota

## Abstract

**Background:** Plant breeders seek to develop cultivars with maximal agronomic value. The merit of breeding material is often assessed using numerous, often genetically correlated traits. As intervention on one trait will affect the value of another, breeding decisions should consider the relationships among traits in the context of putative causal structures (i.e., trait networks). With the proliferation of multi-trait genome-wide association studies (MTM-GWAS), we can infer putative genetic signals at the multivariate scale. However, a standard MTM-GWAS does not accommodate the network structure of phenotypes, and therefore does not address how the traits are interrelated.

**Results:** We extended the scope of MTM-GWAS by incorporating trait network structures into GWAS using structural equation models (SEM-GWAS). In this network GWAS model, the learned structure is used to define a set of explanatory variables that describe how other phenotypes may act on the focal trait. A salient feature of SEM-GWAS is that it can partition the total single nucleotide polymorphism (SNP) effects into direct and indirect effects. Here, we illustrate the utility of SEM-GWAS using a digital metric for shoot biomass, root biomass, water use, and water use efficiency in rice.

**Conclusions:** We found that SNPs impacted water use efficiency directly as well as indirectly through shoot biomass and root biomass. In addition, SEM-GWAS partitioned significant SNP effects influencing water use efficiency into direct and indirect effects as a function of the other traits, providing further biological insights. These results suggest that the use of SEM may enhance our understanding of complex relationships among agronomic traits.

## Introduction

Elite cultivars are the result of generations of targeted selection for multiple characteristics. In many cases, plant and animal breeders alike seek to improve many, often correlated, phenotypes simultaneously. Thus, breeders must consider the interaction between traits during selection. For instance, genetic selection for one trait may increase or decrease the expression of another trait, depending on the genetic correlation between the two. While consideration of the genetic correlation between traits is essential in this respect, modeling recursive interactions between phenotypes provides important insights for developing breeding and management strategies for crops that cannot be realized with conventional multivariate approaches alone. In particular, inferring the structure of trait networks from observational data is critical for our understanding of the interdependence of multiple phenotypes [1–3].

Genome-wide association studies (GWAS) have become increasingly popular approaches for the elucidation of the genetic basis of economically important traits. They have been successful in identifying single nucleotide polymorphisms (SNPs) associated with a wide spectrum of phenotypes, including yield, abiotic and biotic stresses, and plant morphological traits [4]. For many studies, multiple, often correlated, traits are recorded on the same material, and association mapping is performed for each trait independently. While such approaches may yield powerful, biologically meaningful results, they fail to adequately capture the genetic interdependancy among traits and impose limitations on elucidating the genetic mechanisms underlying a complex system of traits. When multiple phenotypes possess correlated structures, multi-trait GWAS (MTM-GWAS), which is the application of mutli-trait models (MTM) [5] to GWAS, is the standard approach. The rationale behind this is to leverage genetic correlations among phenotypes to increase statistical power for the detection of quantitative trait loci, particularly for traits that have low heritability or are scarcely recorded.

While MTM-GWAS is a powerful approach to capture the genetic correlations between traits for genetic inference, it fails to address how the traits are interrelated, or elucidate the mechanisms that give rise to the observed correlation. The early work of Sewall Wright sought to infer causative relations between correlated variables through path analysis [6]. This seminal work gave rise to structural equation models (SEM), which assesses the nature and magnitude of direct and indirect effects of multiple interacting variables. Although SEM remains a powerful approach to model the relationships among variables in complex systems, its use has been limited in biology.

Recently, Momen et al. [7] proposed the SEM-GWAS framework by incorporating trait networks and SNPs into MTM-GWAS through SEM [6, 8]. In contrast to standard multivariate statistical techniques, the SEM framework opens up a multivariate modeling strategy that accounts for recursive (an effect from one phenotype is passed onto another phenotype) and simultaneous (reciprocal) structures among its variables [9, 10]. Momen et al. [7] showed that SEM-GWAS can supplement MTM-GWAS, and is capable of partitioning the source of the SNP effects into direct and indirect effects, which helps to provide a better understanding of the relevant biological mechanisms. In contrast, MTM-GWAS, which does not take the network structure between phenotypes into account, estimates overall SNP effects that are mediated by other phenotypes, and combines direct and indirect SNP effects.

Current climate projections predict an increase in the incidence of drought events and elevated temperatures throughout the growing season [11]. These elevated temperatures will drive higher evapotranspirational demands, and combined with the increased unpredictability of precipitation events, will increase the frequency and intensity of drought, thus impacting crop productivity [12–16]. To mitigate the effects of climate change on agricultural productivity, the development of drought-tolerant cultivars is important for increasing climate resilience in agriculture. However, progress towards this goal is often hindered by the inherent complexity of traits such as drought tolerance [17–20]. The ability to mitigate yield losses under limited water conditions involves a suite of morphological and physiological traits [20]. Among these is the ability to access available water and utilize it for growth. Thus, studying traits associated with water capture (e.g., root biomass and architecture) and utilization (e.g., water-use efficiency) is essential. However, of equal importance is a robust statistical framework that allows these complex traits to be analyzed jointly and network relationships among traits to be inferred for efficient incorporation of these traits into breeding programs.

In this study, we applied SEM-GWAS and MTM-GWAS to incorporate the trait network structures related to shoot and root biomass and to drought responses in rice (*Oryza sativa* L.) from a graphical modeling perspective. Graphical modeling offers statistical inferences regarding complex associations among multivariate phenotypes. Plant biomass and drought stress responses are interconnected through physiological pathways that may be related to each other, requiring the specification of recursive effects using SEM. We combined GWAS with two graphical modeling approaches: a Bayesian network was used to infer how each SNP affects a focal phenotype directly or indirectly through other phenotypes, and SEM was applied to represent the interrelationships among SNPs and multiple phenotypes in the form of equations and path diagrams.

## Materials and Methods

### Experimental data set

The plant material used in our analysis consisted of a rice diversity panel of *n* = 341 inbred accessions of *O. sativa* that originate from diverse geographical regions and are expected to capture much of the genetic diversity within cultivated rice [21]. All lines were genotyped with 700,000 SNPs using the high-density rice array from Affymetrix (Santa Clara, CA, USA) such that there was approximately 1 SNP every 0.54Kb across the rice genome [21, 22]. We used PLINK v1.9 software [23] to remove SNPs with a call rate ≤ 0.95 and a minor allele frequency ≤ 0.05. Missing genotypes were imputed using Beagle software version 3.3.2 [24]. Finally, 411,066 SNPs were retained for further analysis.

### Phenotypic data

We analyzed four biologically important traits for drought responses in rice: projected shoot area (PSA), root biomass (RB), water use (WU), and water use efficiency (WUE). These phenotypes are derived from two previous work [25, 26]. The aim of the first study was to evaluate the effects of drought on shoot growth [26]. Here, the diversity panel was phenotyped using an automated phenotyping platform in Adelaide, SA, Australia. This new phenotyping technology enabled us to produce high-resolution spatial and temporal image-derived phenotypes, which can be used to capture dynamic growth, development, and stress responses [27–30].

The plants were phenotyped over a period of 20 days, starting at 13 days after they were transplanted into soil and ending at 33 days. Each day, the plants were watered to a specific target weight to ensure the soil was completely saturated. The plants were then imaged from three angles (two side views and a top view image). These images were processed to remove all background objects, leaving just pixels for the green shoot tissue. We summed the pixels from each image to obtain an estimate of the shoot biomass. We refer to this metric as PSA. With this system, we also obtained the weights, prior to watering and after watering, for each pot on each day. From this data, we estimated the amount of water that is used by each plant. WU was calculated as Pot Weight_(*r*−1)_ − Pot Weight_(*r*)_, where *r* is time, and WUE is the ratio of PSA to WU. Although this data has not yet been published, a description of the phenotyping system and insight into the experimental design can be found in Campbell et al. [29].

The aim of the second study was to assess salinity tolerance in the rice diversity panel. The plants were grown in a hydroponic system in a greenhouse. Salt stress was imposed for two weeks, and destructive phenotyping performed at 28 days after transplantation. A number of traits were recorded, including RB. The experimental design of this study is fully described in Campbell et al. [25]. All the aforementioned phenotypes were measured under control conditions. The 15th day of imaging was selected for analysis of PSA, WU, and WUE, which is equivalent to 28 days after transplantation, so it matched the age at which RB was recorded. For both studies, best linear unbiased estimates were computed for each accession prior to downstream analyses. For RB, the details of the model are discussed in Campbell et al. [25]. Briefly, a linear model was fitted using the PROC-GLM procedure in SAS that accounted for time of the year, replication, and block effects. For traits derived from high-throughput phenotyping, the linear model included a fixed term for the effect of the experiment and a fixed term for replication nested within experiment.

### Multi-trait genomic best linear unbiased prediction

A Bayesian multi-trait genomic best linear unbiased prediction (MT-GBLUP) model was used for four traits to obtain posterior means of genetic values as inputs for inferring a trait network.

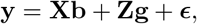

where **y** is the vector observations for *t* = 4 traits, **X** is the incidence matrix of covariates, **b** is the vector of covariate effects, **Z** is the incidence matrix relating accessions with additive genetic effects, **g** is the vector of additive genetic effects, and ***ϵ*** is the vector of residuals. The incident matrix **X** only included intercepts for the four traits examined in this study. Under the infinitesimal model of inheritance, the **g** and ***ϵ*** were assumed to follow a multivariate Gaussian distribution 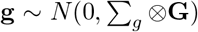 and 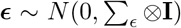, respectively, where **G** is the *n* × *n* genomic relationship matrix for genetic effects, **I** is the identity matrix for residuals, ∑_*g*_ and ∑_*ϵ*_, are the *t* × *t* variance-covariance matrices of genetic effects and residuals, respectively, and ⊗ denotes the Kronecker product. The **G** matrix was computed as 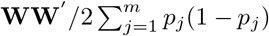, where **W** is the centered marker incidence matrix taking values of 0 − 2*p_j_* for zero copies of the reference allele, 1 − 2*p_j_* for one copy of the reference allele, and 2 − 2*p_j_* for two copies of the reference allele [31]. Here, *p_j_* is the allele frequency at SNP *j* = 1,⋯, *m*. We assigned flat priors for the intercept and the vector of fixed effects. The vectors of random additive genetic effects and residual effects were assigned independent multivariate normal priors with null mean and inverse Wishart distributions for the covariance matrices.

A Markov chain Monte Carlo (MCMC) approach based on Gibbs sampler was used to explore posterior distributions. We used a burn-in of 25,000 MCMC samples followed by an additional 150,000 MCMC samples. The MCMC samples were thinned by a factor of two, resulting in 75,000 MCMC samples for inference. Posterior means were then calculated for estimating model parameters. The MTM R package was used to fit the above regression model (https://github.com/QuantGen/MTM).

### Learning structures using Bayesian network

Networks or graphs can be used to model interactions. Bayesian networks describe conditional independence relationships among multivariate phenotypes. Each phenotype is connected by an edge to another phenotype if they directly affect each other given the rest of the phenotypes, whereas the absence of edge implies conditional independence given the rest of phenotypes. Several algorithms have been proposed to infer plausible structures in Bayesian networks, assuming independence among the realization of random variables [32]. The estimated genetic values from MT-GBLUP were used as inputs, and we applied the Hill Climbing (HC) algorithm from the score-based structure learning category to infer the network structure among the four traits examined in this study [33]. We selected this algorithm because it was suggested in a recent study, [34], which showed that the score-based algorithms performed better for the construction of networks than constraint-based counterparts. The bnlearn R package was used to learn the Bayesian trait network throughout this analysis with mutual information as the test, and the statistically significant level set at *α* = 0.01 [32]. We computed the Bayesian information criterion (BIC) score of a network and estimated the strength and uncertainty of direction of each edge probabilistically by bootstrapping [35]. In addition, the strength of the edge was assessed by computing the change in the BIC score when that particular edge was removed from the network, while keeping the rest of the network intact.

### Multi-trait GWAS

We used the following MTM-GWAS that does not account for the inferred network structure by extending the single-trait GWAS counterpart of Kennedy et al. [36] and Yu et al. [37]. For ease of presentation, it is assumed that each phenotype has null mean.

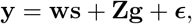

where **w** is the *j*th SNP being tested, **s** represents the vector of fixed *j*th SNP effect, and **g** is the vector of additive polygenic effect. The aforementioned variance-covariance structures were assumed for **g** and ***ϵ***. The MTM-GWAS was fitted individually for each SNP, where the output is a vector of marker effect estimates for each trait, i.e. **ŝ** = [*ŝ*_PSA_, *ŝ*_RB_, *ŝ*_WU_, *ŝ*_WUE_].

### Structural equation model for GWAS

A structural equation model is capable of conveying directed network relationships among multivariate phenotypes involving recursive effects. The SEM described in Gianola and Sorensen [38] in the context of linear mixed models was extended for GWAS, according to [7].

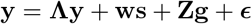

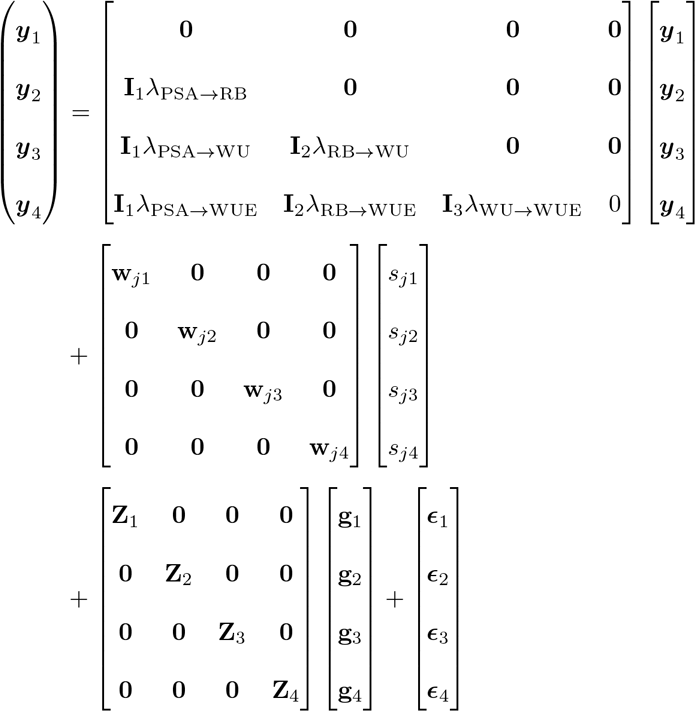

where **I** is the identity matrix, **Λ** is the lower triangular matrix of regression coefficients or structural coefficients based on the learned network structure from the Bayesian network, and the other terms are as defined earlier.

Note that the structural coefficients **Λ** determine that the phenotypes which appear in the left-hand side also appear in the right-hand side, and represent the edge effect size from phenotype to phenotype in Bayesian networks. If all elements of **Λ** are equal to 0, then this model is equivalent to MTM-GWAS. Gianola and Sorensen [38] showed that the reduction and re-parameterization of a SEM mixed model can yield the same joint probability distribution of observation as MTM, suggesting that the expected likelihoods of MTM and SEM are the same [39]. For example, we can rewrite the SEM-GWAS model as

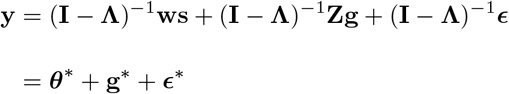

where Var(**g**\*) ~ (**I** − **Λ**)^-1^**G**(**I** − **Λ**)′^-1^ and Var(***ϵ***\*) ~ (**I** − **Λ**)^−1^**R**(**I** − **Λ**)′^-1^. This transformation changes SEM-GWAS into MTM-GWAS, which ignores the network relationships among traits [38, 39]. However, Valente et al. [40] stated that SEM allows for the prediction of the effects of external interventions, which can be useful for making selection decisions that are not possible with MTM. We used SNP Snappy software to perform MTM-GWAS and SEM-GWAS [41]. To identify candidate SNPs that may explain direct (in the absence of mediation by other traits) and indirect (with intervention and mediation by other traits) effects for each trait, the SNPs from MTM-GWAS were ranked according to *p*-values for each trait. The top 50 SNPs were then selected, and marker effects were decomposed into direct and indirect effects using SEM-GWAS. Since WU and WUE were the only two traits to have indirect effects, we focused on these traits for downstream analysis with SEM-GWAS.

## Results

### Trait correlations and network structure

Multi-phenotypes were split into genetic values and residuals by fitting the MT-GBLUP. The estimates of genomic and residual correlations among the four traits measured in this study are shown in Table 1. Correlations between all traits ranged from 0.48 to 0.92 for genomics and –0.13 to 0.83 for residuals. The estimated genomic correlations can arise from pleiotropy or linkage disequilibrium (LD). Although pleiotropy is the most durable and stable source of genetic correlations, LD is considered to be less important than pleiotropy because alleles at two linked loci may become non-randomly associated by chance and be distorted through recombination [42, 43].

**Table 1:** Genomic (upper triangular), residual (lower triangular) correlations and genomic heritablities (diagonals) of four traits in the rice with posterior standard deviations in parentheses. Projected shoot area (PSA), root biomass (RB), water use (WU), and water use efficiency (WUE).

|  | PSA | RB | WU | WUE |
| --- | --- | --- | --- | --- |
| PSA | 0.677 (0.092) | 0.515 (0.102) | 0.846 (0.043) | 0.920 (0.018) |
| RB | 0.030 (0.218) | 0.733 (0.083) | 0.479 (0.114) | 0.517 (0.107) |
| WU | 0.443 (0.152) | -0.134 (0.216) | 0.643(0.097) | 0.744 (0.076) |
| WUE | 0.829 (0.052) | 0.111 (0.195) | 0.106 (0.182) | 0.576 (0.092) |

We postulated that the learned networks can provide a deeper insight into relationships among traits than simple correlations or covariances. Figure 1 shows a network structure inferred using the hill-climb (HC) algorithm. This is a fully recursive structure because there is at least one incoming or outgoing edge for each node. Unlike the MTM-GWAS model, the inferred graph structure explains how the phenotypes may be related to each other either directly or indirectly mediated by one or more variables. We found a direct dependency between PSA and WUE. A direct connection was also found between RB and WUE, and PSA and WU.

**Figure 1:**
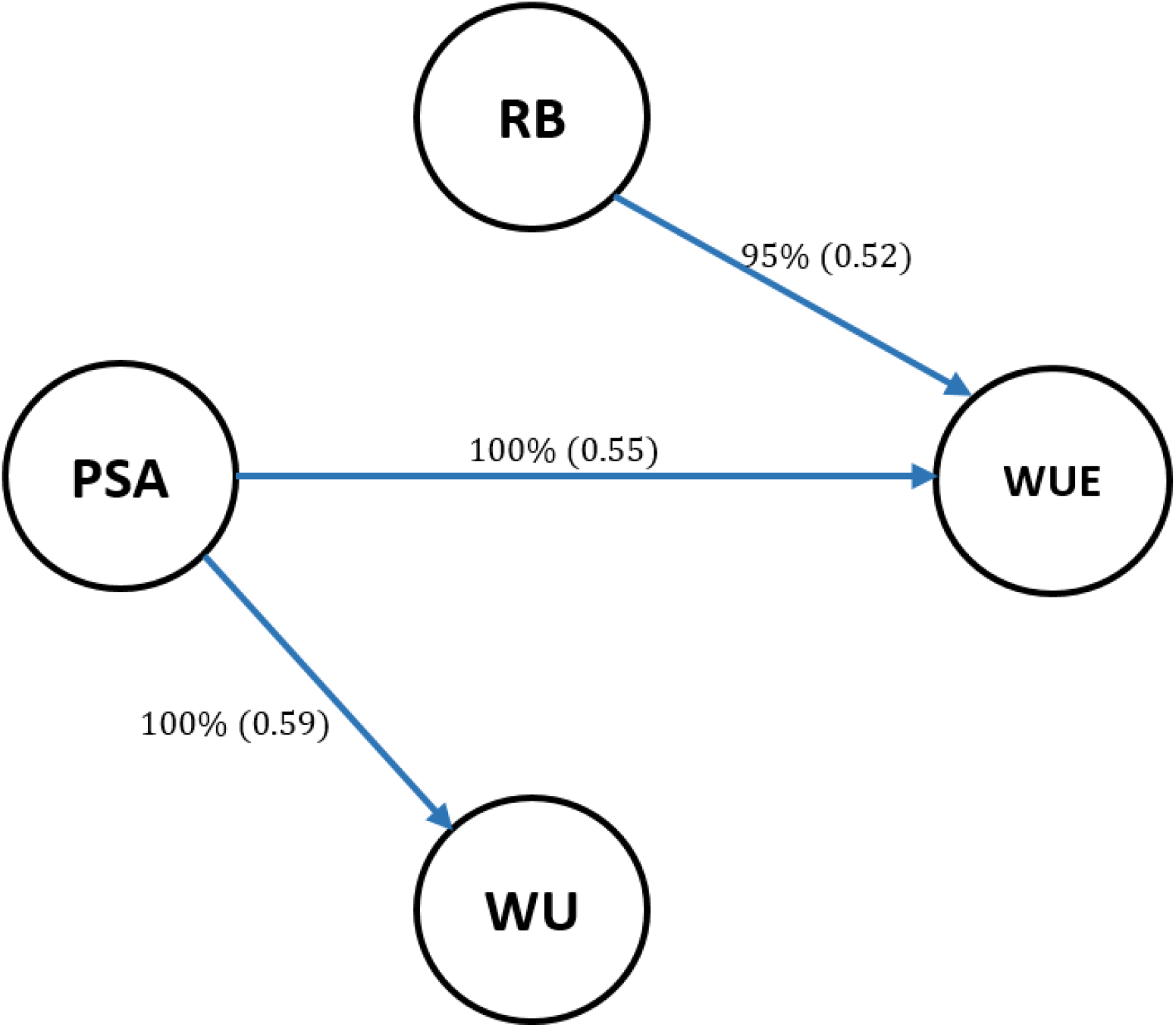
Scheme of inferred network structure using the Hill-Climbing (HC) algorithm, with 0.85, threshold; the minimum strength required for an arc to be included in the network. Structure learning test was performed with 2,500 bootstrap samples with mutual information as the test statistic with a significance level at *α* = 0.01. Labels of the edges refer to the strength and direction (parenthesis) which measure the confidence of the directed edge. The strength indicates the frequency of the edge is present and the direction measures the frequency of the direction conditioned on the presence of edge. PSA: projected shoot area; RB: root biomass; WU: water use; WUE: water use efficiency.

Measuring the strength of probabilistic dependence for each arc is crucial in Bayesian network learning [35]. As shown in Figure 1, the strength of each arc was assessed with 2,500 bootstrap samples with a significance level at *α* = 0.01. The labels on the edges indicate the proportion of bootstrap samples supporting the presence of the edge and the proportion supporting the direction of the edges are provided in parentheses. Learned structures were averaged with a strength threshold of 85% or higher to produce a more robust network structure. Edges that did not meet this threshold were removed from the networks. In addition, we used BIC as goodness-of-fit statistics measuring how well the paths mirror the dependence structure of the data (Table 2). The BIC assign higher scores to any path that fit the data better. The BIC score reports the importance of each arc by its removal from the learned structure. We found that removing PSA → WUE resulted in the largest decrease in the BIC score, suggesting that this path is playing the most important role in the network structure. This was followed by PSA → WU and RB → WUE.

**Table 2:** Bayesian information criterion (BIC) for the network learned using the hill-climbing (HC) algorithm. BIC denote BIC scores for pairs of nodes and reports the change in the score caused by an arc removal relative to the entire network score. Projected shoot area (PSA), root biomass (RB), water use (WU), and water use efficiency (WUE).

| Algorithm | from | to | BIC |
| --- | --- | --- | --- |
| HC | PSA | WU | -427.956 |
|  | PSA | WUE | -488.787 |
|  | RB | WUE | -3.327 |

### Structural equation coefficients

The inferred Bayesian network among PSA, RB, WU, and WUE in Figure 1 was modeled using a set of structural equations to estimate SEM parameters and SNP effects, as shown in Figure 2, which can be statistically expressed as

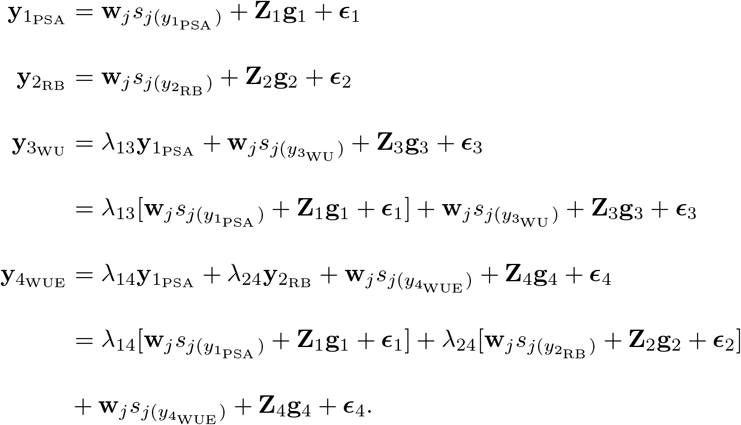

**Figure 2:**
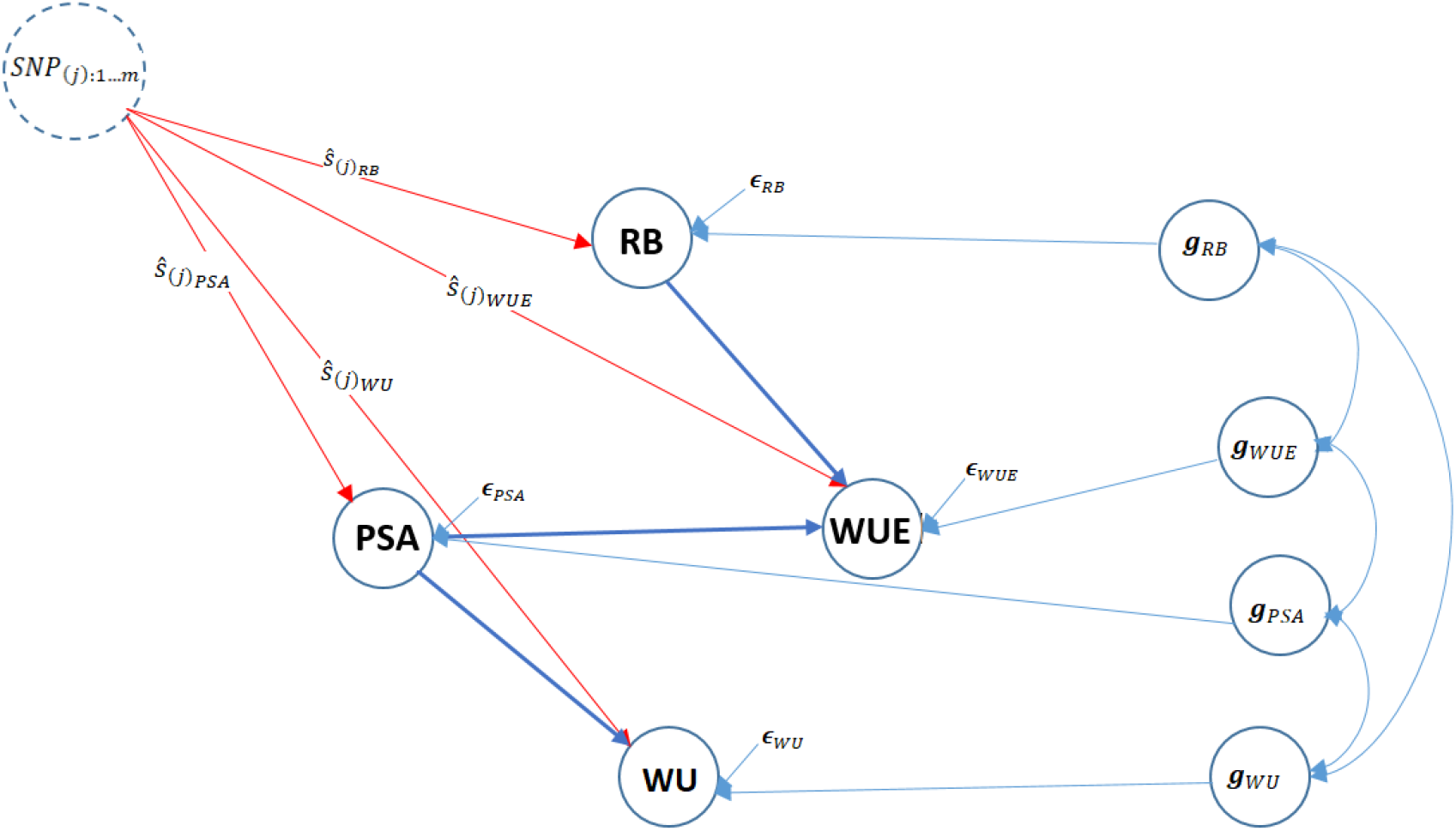
Pictorial representation of trait network and SNP effects (*ŝ*) using the structural equation model for four traits. Unidirectional arrows indicate the direction of effects and bidirectional arrows represent genetic correlations (*g*) among phenotypes. PSA: projected shoot area; RB: root biomass; WU: water use; WUE: water use efficiency; *ϵ*: residual.

The corresponding estimated **Λ** matrix is

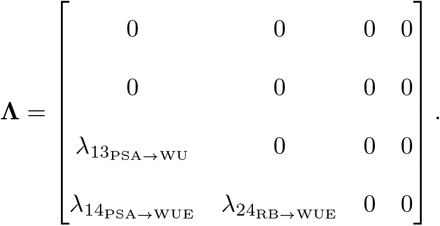

Table 3 presents the magnitude of estimated structural path coefficients: λ_13_, λ_14_, and λ_24_ for PSA on WU, PSA on WUE, and RB on WUE, respectively. The structural coefficients (λ_*ii*′_) describe the rate of change of trait *i* with respect to trait *i*′. The largest magnitude of the structural coefficient was 0.963, which was estimated for PSA→WUE, whereas the lowest was 0.045, which was estimated for RB→WUE.

**Table 3:** Structural coefficients (λ) estimates derived from the structural equation models. Projected shoot area (PSA), root biomass (RB), water use (WU), and water use efficiency (WUE).

| Path | $\lambda$ | Structural coefficient |
| --- | --- | --- |
| PSA → WU | $\lambda_{13}$ | 0.761 |
| PSA → WUE | $\lambda_{14}$ | 0.963 |
| RB → WUE | $\lambda_{24}$ | 0.045 |

### Interpretation of SNP effects

We implemented SEM-GWAS as an extension of the MTM-GWAS method for analysis of the joint genetic architecture of the four measured traits, to partition SNP effects into direct and indirect [44]. The results of the decomposition of SNP effects are discussed for each trait separately below. Because the network only revealed indirect effects for WU and WUE, we focused on these traits for decomposing marker effects.

#### Projected shoot area (PSA)

Figure 3 shows a Manhattan plot of SNP effects on the PSA. According to the path diagram, there is no intervening trait or any mediator variable for PSA (Figure 2). It is possible that the PSA architecture is only influenced by the direct SNP effects, and is not affected by any other mediators or pathways. Hence, the total effect of *j*th SNP on PSA is equal to its direct effects.

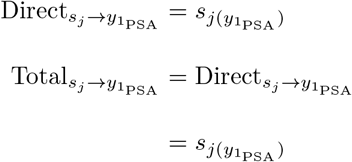

**Figure 3:**
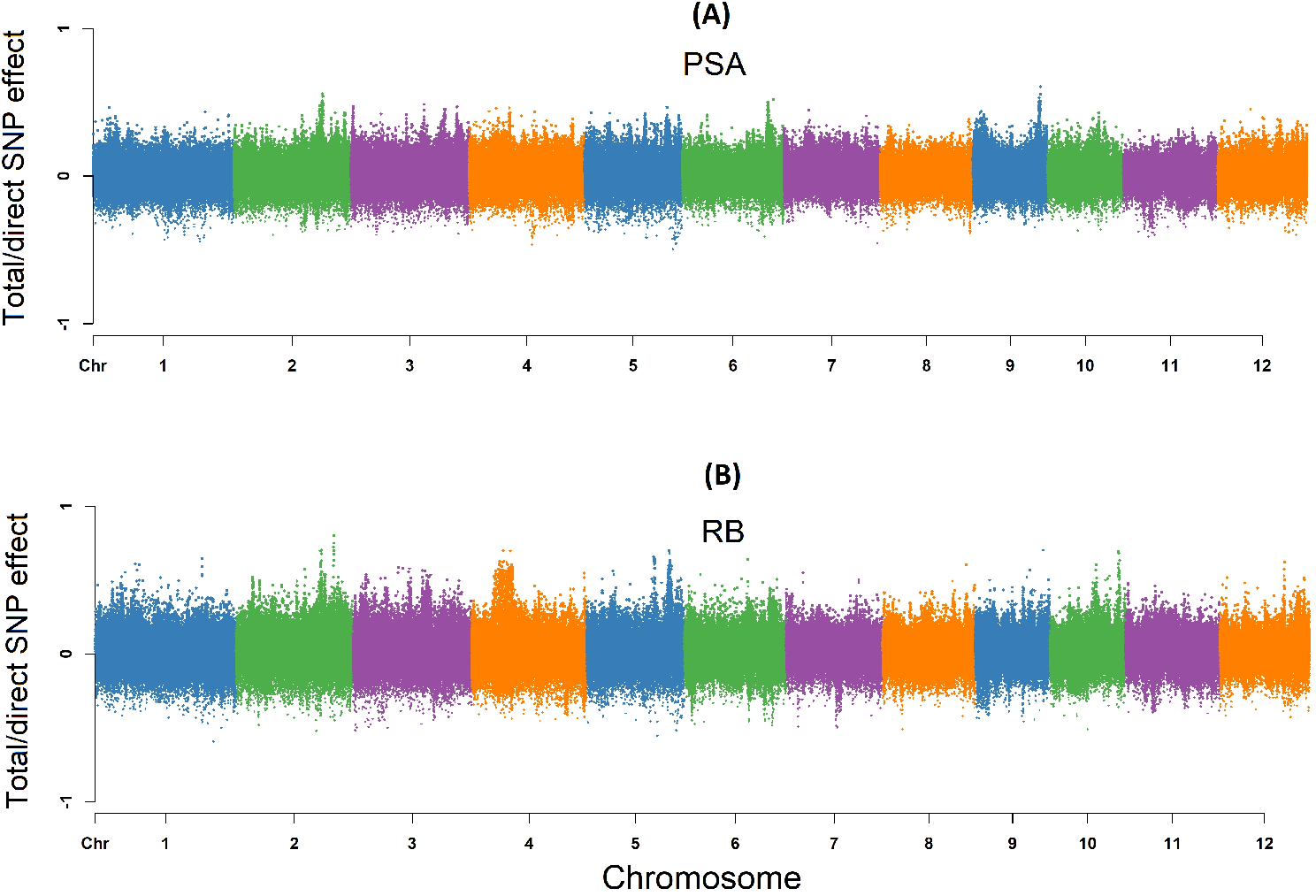
Manhattan plots (total/direct) SNP effects on projected shoot area (PSA) and root biomass (RB) using SEM-GWAS based on the network learned by the hill climbing algorithm. Each point represents a SNP and the height of the SNP represents the extent of its association with PSA and RB.

#### Root biomass (RB)

No incoming edges were detected for RB, resulting in a similar pattern to PSA, which suggests that SNP effects on RB were not mediated by other phenotypes. As shown in Figure 3, a Manhattan plot for RB consists of direct and total effects.

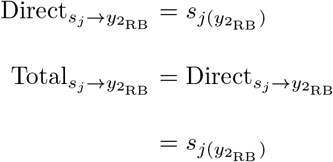

#### Water use (WU)

Based on Figure 2, the total effects for a single SNP can be decomposed into direct effects on WU and indirect effects in which PSA acts as a mediator as WU has a single incoming edge from PSA. Thus, the SNP effect transmitted from PSA contribute to the total SNP effect on WU. Under these conditions, the estimated total SNP effects for WU cannot be simply described as the direct effect of a given SNP, since the indirect effect of PSA must also be considered. This is different from MTM-GWAS, which does not distinguish between the effects mediated by mediator phenotypes, and only captures the overall SNP effects. Here it should be noted that the extent of SNP effects from PSA on WU are controlled by the structural equation coefficients λ_13_. Figure 4 shows a Manhattan plot of SNP effects on WU.

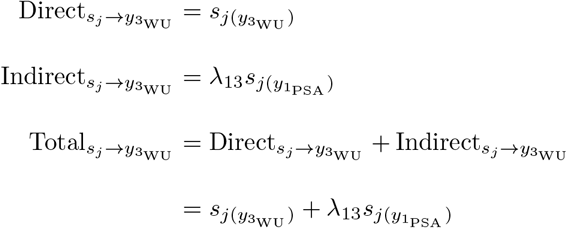

**Figure 4:**
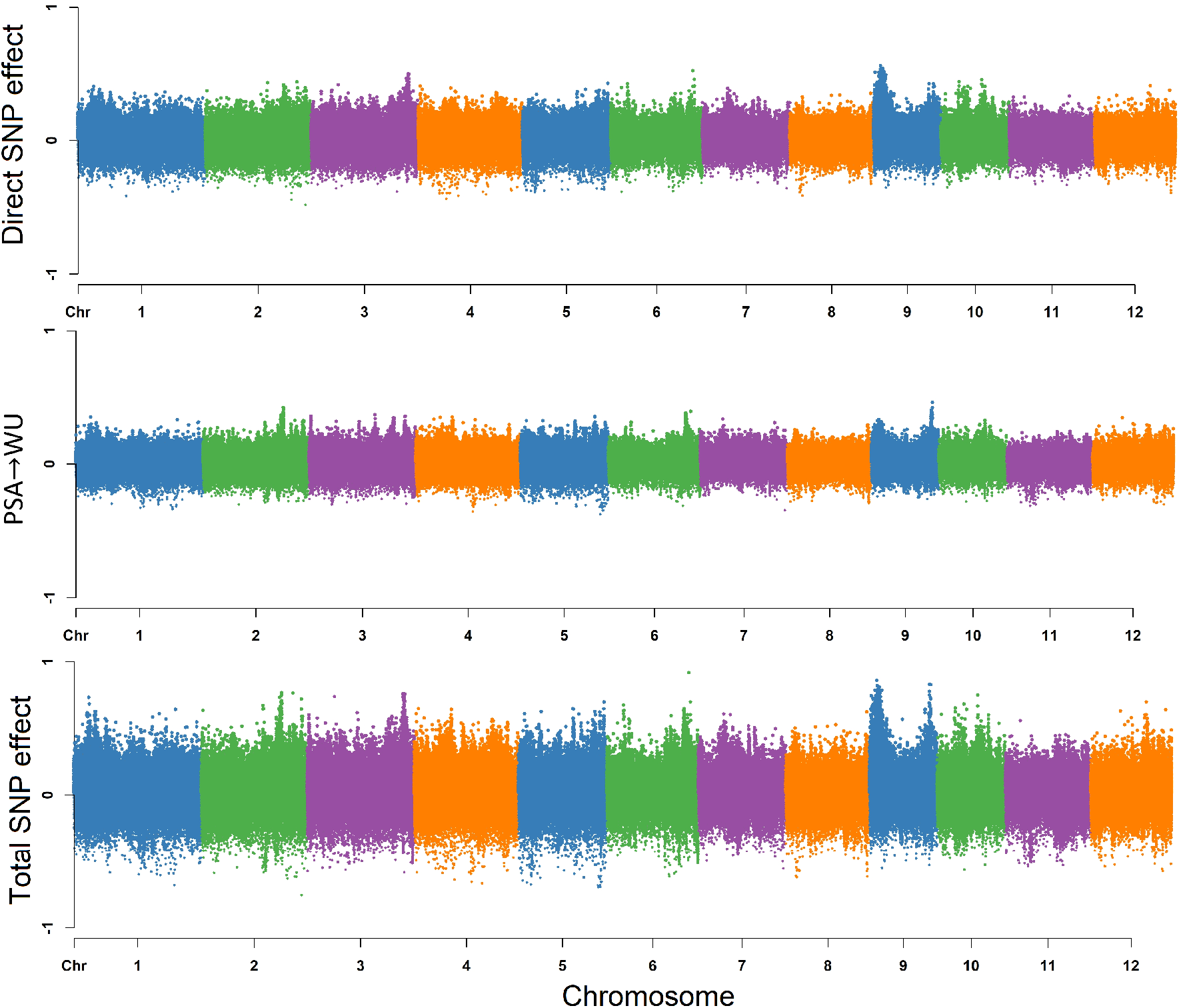
Manhattan plot of direct (affecting each trait without any mediation), indirect (mediated by other phenotypes), and total (sum of all direct and indirect) SNP effects on water use (WU) using SEM-GWAS based on the network learned by the hill climbing algorithm. Each point represents a SNP and the height of the SNP represents the extent of its association with WU.

#### Water use efficiency (WUE)

The overall SNP effects for WUE can be partitioned into one direct and two indirect genetic signals (Figure 2). WU and WUE are the traits that do not have any outgoing path to other traits. According to Figure 5, the extents of the SNP effects among the two indirect paths were 1) RB → WUE, and 2) PSA→WUE in increasing order. We found that the SNP effect transmitted through RB had the smallest effect on WUE, suggesting that modifying the size of the QTL effect for RB may not have a noticeable effect on WUE, whereas a change in PSA may have a noticeable effect on WUE. The magnitude of the relationship between RB and WUE is proportional to the product of structural coefficients λ_24_ = 0.045. PSA influenced WUE via a single indirect path, and strongly depends on the structural coefficient λ_14_ = 0.963 for PSA → WUE. Collectively these results suggest that WUE can be influenced by selection on PSA.

**Figure 5:**
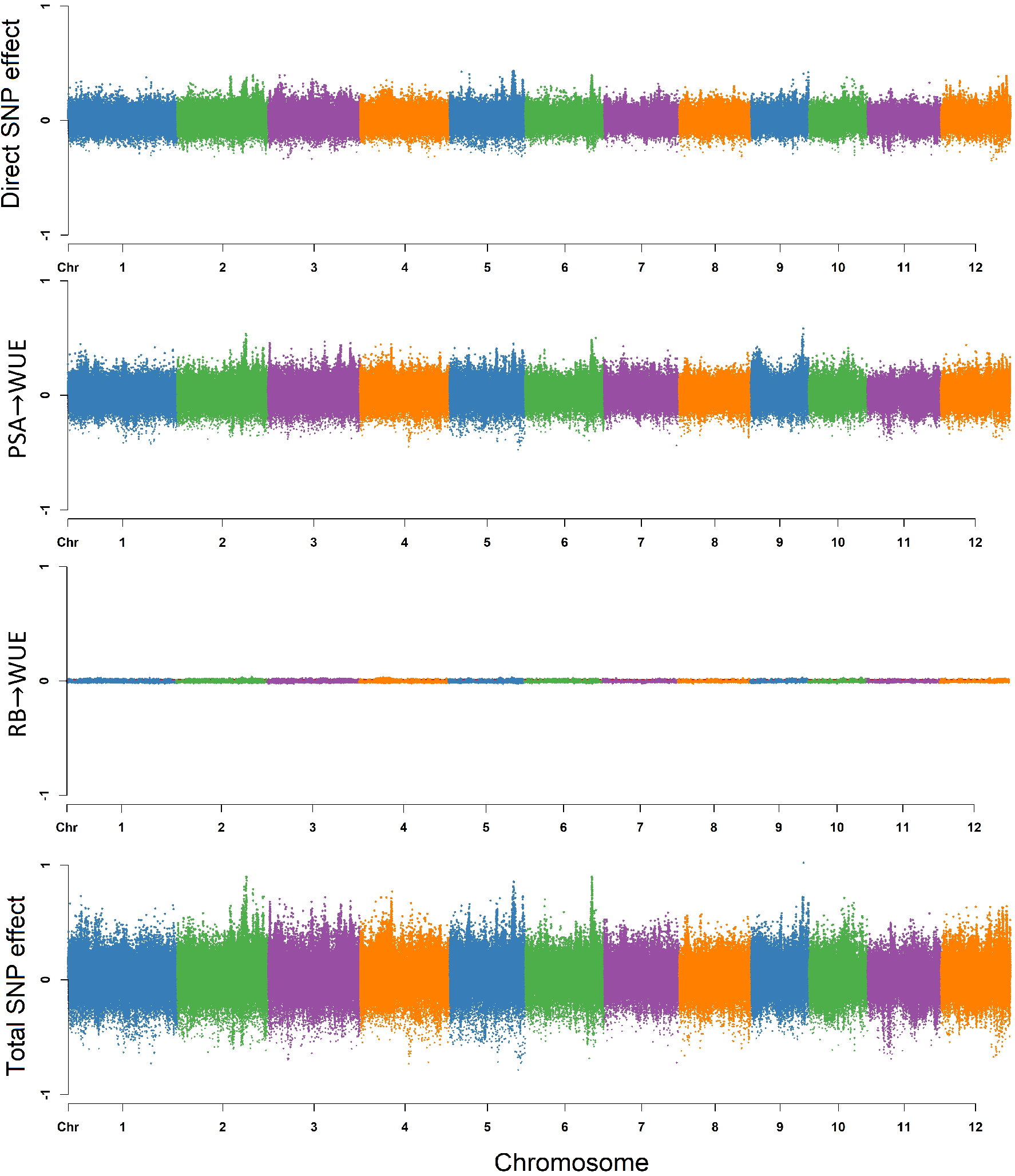
Manhattan plot of direct (affecting each trait without any mediation), indirect (mediated by other phenotypes), and total (sum of all direct and indirect) SNP effects on water use efficienty (WUE) using SEM-GWAS based on the network learned by the hill climbing algorithm. Each point represents a SNP and the height of the SNP represents the extent of its association with WUE.

The direct and indirect effects are summarized with the following equation:

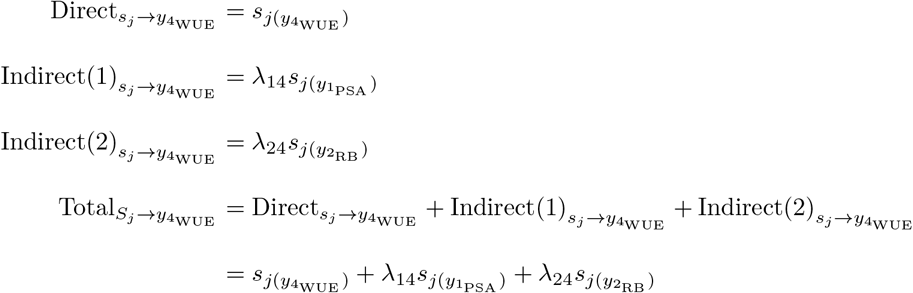

### Leveraging SEM-GWAS to decompose pleiotropic QTL

Pleiotropy can be simply defined as a gene that has an effect on multiple traits, however understanding how the gene acts on multiple traits is a challenge. The advantage of SEM-GWAS is that it can be used to understand how a QTL acts on multiple interrelated traits. Thus, it can be used to decompose pleiotropic QTL effects into direct and indirect effects, and understand how a given QTL acts on multiple traits. We next sought to identify QTL with pleiotropic effects and elucidate how the QTL acts on the traits. To this end, we ranked SNPs from MTM-GWAS based on p-values to select the top 50 SNPs for each trait and used SEM-GWAS to elucidate how marker effects were partitioned among traits. Since the inferred network revealed indirect effects for only WU and WUE, downstream analyses were focused on these two traits.

Top SNPs for WU and WUE showed very different patterns of pleiotropy. For WU, the direct SNP effect size was on average 57% higher than the indirect SNP effect size coming from PSA, indicating that the total SNP effects from WU are driven largely by genetic effects acting directly on WU rather than indirectly thought PSA. However for WUE, direct SNP effects on WUE had a much smaller contribution to total SNP effects compared to indirect effects from PSA. For instance, comparisons between direct SNP effect on WUE and indirect effects from PSA on WUE showed that direct effects were, on average, 16% lower than indirect effects. While indirect contributions from RB on total SNP effects were minimal, with indirect effects from RB on WUE showing an approximately 30 fold lower effect than direct effects on WUE. Thus, for many loci associated with WUE, the total effects may be driven largely by the marker’s effect on PSA rather than WUE directly. These patterns may be due to the very high genomic correlation between PSA and WUE.

While most of the top SNPs from MTM for WU showed larger direct effects on WU compared to indirect effects through PSA, several loci were identified where direct effects were nearly equal to indirect effects. For instance, the direct effect on WU for SNP-4.30279060. was −0.272, while the indirect effect through PSA was −0.268. Moreover, this SNP was the second most significant SNP associated with PSA from MTM-GWAS. The effects of this SNP on both PSA and WU is apparent in Figure 6. Individuals with the “2” allele had considerably lower shoot biomass and lower water use than those with the “0” allele. Conversely, SNPs with small indirect effects on WU through PSA relative to direct effects on WU were ranked much lower for MTM-GWAS for PSA. The SNP-10.2860531. had considerably smaller indirect effect on WU through PSA relative to the direct effect on WU (−0.124 and −0.327, respectively) on WU, and was ranked 17,902 for PSA from MTM-GWAS.

**Figure 6:**
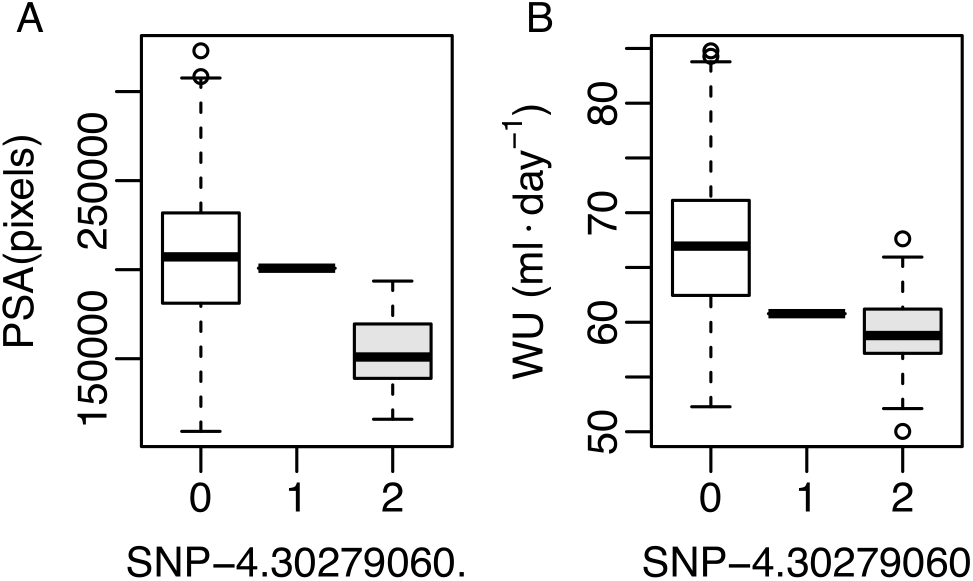
Distribution of projected shoot area (PSA) and Water use (WU) for allelic groups at SNP-4.30279060. PSA values are shown in A, while water use values are shown in B. The *x*-axis shows allele counts at SNP-4.30279060, where 0, 1 and 2 indicate accessions that are homozygous for the reference allele, heterozygous, and homozygous for the alternative allele.

To further examine the putative biological effects of these loci, we next sought to identify candidate genes near SNPs of interest. To this end, we extracted genes within a 200kb window of each SNP. Several no-table genes were identified that have reported role in regulating plant growth and development, hormone biosynthesis or abiotic stress responses. For instance, a gene encoding a gibberellic acid catabolic protein (*GA2ox7*) was identified approximately 3.5 kb downstream from a SNP (SNP-1.5964363.) associated with WUE through MTM-GWAS (Table 4) [45, 46]. Interestingly, SEM-GWAS revealed that indirect effect from PSA on WUE was approximately 57% greater than direct effects on WUE (*ŝ* = −0.335 and −0.213, respectively). In addition to *OsGA2OX7*, we identified a second gene, *OVP1*, that was associated with WUE. *OVP1* is known to influence abiotic stress responses in rice, as well as growth and development in Arabidopsis [47–49]. Like *OsGA2OX7*, the SNP closest to OVP1 showed larger indirect effects from PSA on WUE than direct effects (*ŝ* = 0.430 and 0.344, respectively).

**Table 4:** Candidate genes for water use efficiency (WUE) identified through SEM-GWAS. Chr: chromosome; BP: gene position in base pairs; GA: gibberellic acid

| Gene ID | Chr | BP | SNP | Rice Annotation | Putative Function | Reference |
| --- | --- | --- | --- | --- | --- | --- |
| LOC_Os01g11150 | 1 | 5,968,819 | SNP-1.5964363. | *GA2OX7* | GA catabolism | [45] |
| LOC_Os01g11054 | 1 | 5,899,555 | SNP-1.5964363. | *OsPPC4* | Growth, $\text{NH}_4^+$ assimilation | [63] |
| LOC_Os06g43660 | 6 | 26,272,897 | SNP-6.26293126. | *OVP1* | Plant growth | [47] |

Several notable genes were identified for WU that have reported roles in regulating plant development and drought tolerance (Table 5). For instance, a gene encoding a lipid transfer protein (*OsDIL1*) was identified approximately 24kb upstream of a SNP associated (SNP-10.2860531.) with WU through MTM-GWAS. Guo et al. [50] showed that plants overexpressing *OsDIL1* were more tolerant to drought stress during the vegetative stage. Examination of the SNP effects through SEM-GWAS revealed that the total SNP effect from MTM-GWAS was primarily driven by direct effect on WU rather than indirect effects on WU through PSA (*ŝ* = −0.327 and −0.124, respectively). In contrast to the locus harboring *OsDIL1*, a region on chromosome 4 was identified that harbored a gene known to regulate growth and development in rice, *MPR25* [51].

**Table 5:** Candidate genes for water use (WU) identified through SEM-GWAS. Chr: chromosome; BP: gene position in base pairs

| Gene ID | Chr | BP | SNP | Rice Annotation | Putative Function | Reference |
| --- | --- | --- | --- | --- | --- | --- |
| LOC_Os01g71990 | 1 | 41,718,016 | SNP-1.41687755. | *P5C* | Proline biosynthesis | [64] |
| LOC_Os04g51350 | 4 | 30,410,105 | SNP-4.30279060. | *MPR25* | Plant development | [51] |
| LOC_Os10g05720 | 10 | 2,885,293 | SNP-10.2860531. | *OsDIL1* | Drought tolerance | [50] |

## Discussion

The relationship between biomass and WU in rice may involve complex network pathways with recursive effects. These network relationships cannot be modeled using a standard MTM-GWAS model. In this study, we incorporated the network structure between four phenotypes, PSA, RB, WU, and WUE, into a multivariate GWAS model using SEM. In GWAS, a distinction between undirected edges and directed edges is crucial, because often biologists and breeders are interested in studying and improving a suite of traits rather than a single trait in isolation. Moreover, intervention on one trait often influences the expression of another [52]. As highlighted in Alwin and Hauser [44], one of the advantages of SEM is that it is capable of splitting the total effects into direct and indirect effects. In regards to genetic studies, SEM enables the researcher to elucidate the underlying mechanism by which an intervention trait may influence phenotypes using a network relationship [53, 54].

Detecting putative causal genes is of considerable interest for determining which traits will be affected by specific loci from a biological perspective, and consequently partitioning the genetic signals according to the paths determined. Although the parameter interpretations of SEM as applied to QTL mapping [55, 56], expression QTL [57], or genetic selection [40] have been actively pursued, the work of Momen et al. [7] marks one of the first studies to account for the level of individual SNP effect in genome-wide SEM analyses. The SEM embeds a flexible framework for performing such network analysis in a GWAS context, and the current study demonstrates its the first application in crops. We assumed that modeling a system of four traits in rice simultaneously may help us to examine the sources of SNP effects in GWAS in greater depth. Therefore, we used two GWAS methodologies that have the ability to embed multiple traits jointly, so that the estimated SNP effects from both models have different meanings. The main difference between SEM-GWAS and MTM-GWAS is that the former includes the relationship between SNPs and measured phenotypes, coupled with relationships that are potentially mediated by other phenotypes (mediator traits). This advances GWAS, and consequently the information obtained from trait networks describing such interrelationships can be used to predict the behavior of complex systems [7]. Although we analyzed the observed phenotypes in the current study, the factor analysis component of SEM can be added to SEM-GWAS by deriving latent factors from multiple phenotypes [e.g., 58, 59]. The inference of a trait network structure was carried out using a Bayesian network, which has applications in genetics ranging from modeling linkage disequilibrium [60] to epistasis [61].

Effective water use and water capture are essential for the growth of plants in arid environments, where water is a limiting factor. These processes are tightly intertwined, and therefore must be studied in a holistic manner. In the current study, we sought to understand the genetic basis of water use, water capture, and growth by examining PSA, RB, WU, and WUE in a diverse panel of rice accessions. The identification of several QTL that affect one or more of these processes highlights the interconnectedness of PSA, RB, WU, and WUE. Water use is a complex trait that is affected by several morphological characteristics (e.g. leaf area, stomatal density, leaf anatomical features, root architecture, anatomy, etc.), and physiological processes (e.g. stomatal aperture) that are greatly influenced by the environment. Thus, any approach that can partition genetic effects for WU among the multiple biological processes that may influence this trait can greatly enhance our understanding of how WU is regulated. Although many of the factors influencing WU were unaccounted for in the current study, the automated phenotyping platform provided an effective means to quantify water use for each plant while simultaneously quantifying shoot biomass. Thus, with these data and the SEM-GWAS framework we can begin to uncouple the complex interrelationship between plant size and water use.

Several QTL were identified for WU through MTM-GWAS. SEM-GWAS revealed that for most loci, the total SNP effect was driven largely by direct effects on WU rather than indirect effects on WU through PSA. In contrast, SEM-GWAS showed that for WUE, total SNP effects were driven largely by effects originating from PSA and acting indirectly on WUE. In the current study, WUE is a composite trait that is defined as the ratio of PSA to WU. The genomic correlation for PSA and WUE was quite high. Although genetic correlation may be due to pleiotropy or linkage disequilibrium, given the definition of WUE the high genetic correlation is likely largely due to the plieotropy [62]. Thus, these two traits are likely controlled by similar QTL, and so it may be very difficult to partition total QTL effect into direct and indirect paths.

Several of the candidate genes associated with loci from MTM-GWAS shed light on the possible biological mechanisms underlying pleiotropic relationships for WU and WUE with PSA. For instance, a SNP located on chromosome 4 was identified for WU and harbored a gene encoding a pentatricopeptide repeat protein (*MPR25*) A closer inspection of this region with SEM-GWAS showed that total SNP effects on WU were largely due to indirect effects originating from PSA. Toda et al. [51] showed that *MPR25* participates in RNA editing and disruption of this gene results in slow growing plants with reduced chlorophyll content. Although considerable work is necessary to determine if *MPR25* underlies natural variation for shoot growth (i.e., PSA) and water use, the presence of this gene near this SNP and the effects of this SNP on PSA and WU present an interesting direction for future studies. In addition to *MPR25*, a second gene was found near a SNP associated with WUE that had a large indirect effect through PSA, *GA2OX7*. The *GA2OX* gene family are involved in the catabolism of the growth promoting hormone gibberellic acid (GA) [45, 46]. GA play important roles in many processes, but are most well known for their role in shaping semi-dwarf rice and wheat cultivars [45, 46]. Modifications in shoot size are likely to influence water use, as larger plants will have greater surface are for evapotranspiration. Thus the presence of this gene within this region on chromosome 1 may explain the larger indirect effects on WUE through PSA compared to the direct effects on WUE.

A deep understanding of the complex relationship between effective water use and water capture, and its impact on plant growth in arid environments, is critical as we continue to develop germplasm that is resilient to climatic variability. As with the significant recent advances in phenomics and remote sensing technologies, future plant breeders will have a new suite of tools to quantify morphological, physiological, and environmental variables at high resolutions. To fully harness these emerging technologies and leverage these multi-dimensional datasets for crop improvement, new analytical approaches must be developed that integrate genomic and phenomic data in a biologically meaningful framework. This study examined multiple phenotypes using a Bayesian network that can serve as potential factors to allow intervention in complex trait GWAS. The SEM-GWAS seems to provide enhanced statistical analysis of MTM-GWAS by accounting for trait network structures.

## Conclusions

We extended the scope of multivariate GWAS by incorporating trait network structures into GWAS using SEM. The main significance of SEM-GWAS is to include the relationship between SNPs and measured phenotypes, coupled with relationships that are potentially mediated by other phenotypes. Using four traits in rice, we showed that SEM-GWAS can partition the total SNP effects into direct and indirect effects, offering new perspectives.

## Declarations

### Acknowledgements

We thank Haipeng Yu for helping with data analysis.

### Availability of data

Genotypic data regarding the rice accessions can be downloaded from the rice diversity panel website (http://www.ricediversity.org/). Phenotypic data used herein are available in Additional File S1.

### Authors’ contributions

MTC and HW designed and conducted the experiments. MM and MTC analyzed the data. MM and GM conceived the idea and wrote the manuscript. MTC and HW discussed results and revised the manuscript. GM supervised and directed the study. All authors read and approved the manuscript.

### Ethics approval and consent to participate

Not applicable.

### Consent for publication

Not applicable.

