## Supplementary Figures for "Utilizing trait networks and structural equation models as tools to interpret multi-trait genome-wide association studies" (from another version of this preprint: version 1, 2019-02-18)

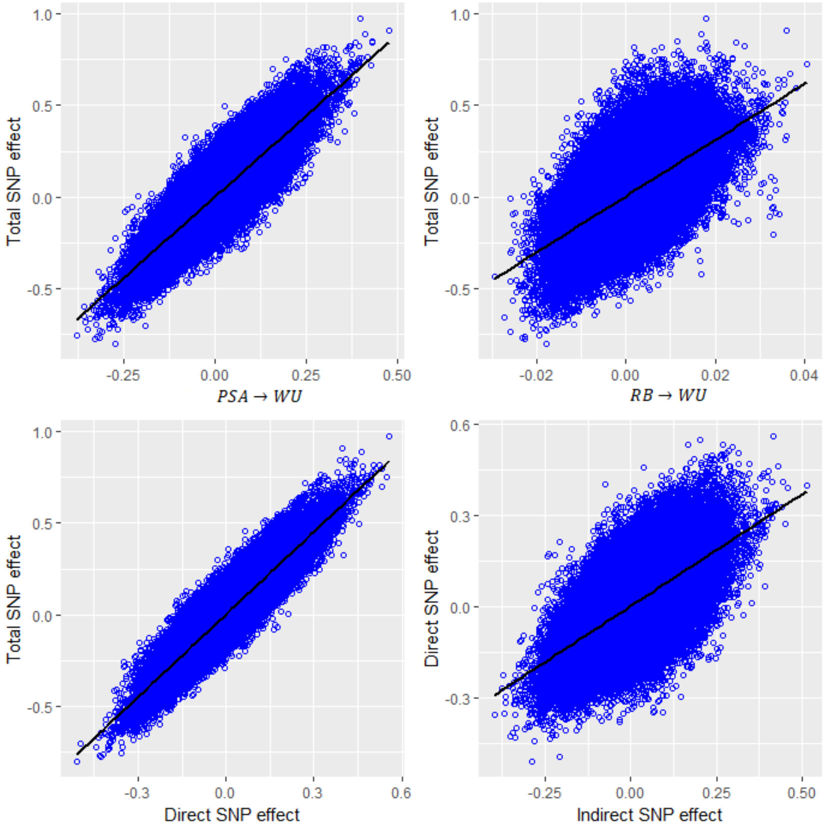


Figure S1: Scatter plots of decomposed SNP effects for water usage (WU). Each point represents the estimated SNP effect for a SNP which affect WU directly or indirectly. PSA: projected shoot area; RB: root biomass; WUE: water use efficiency.

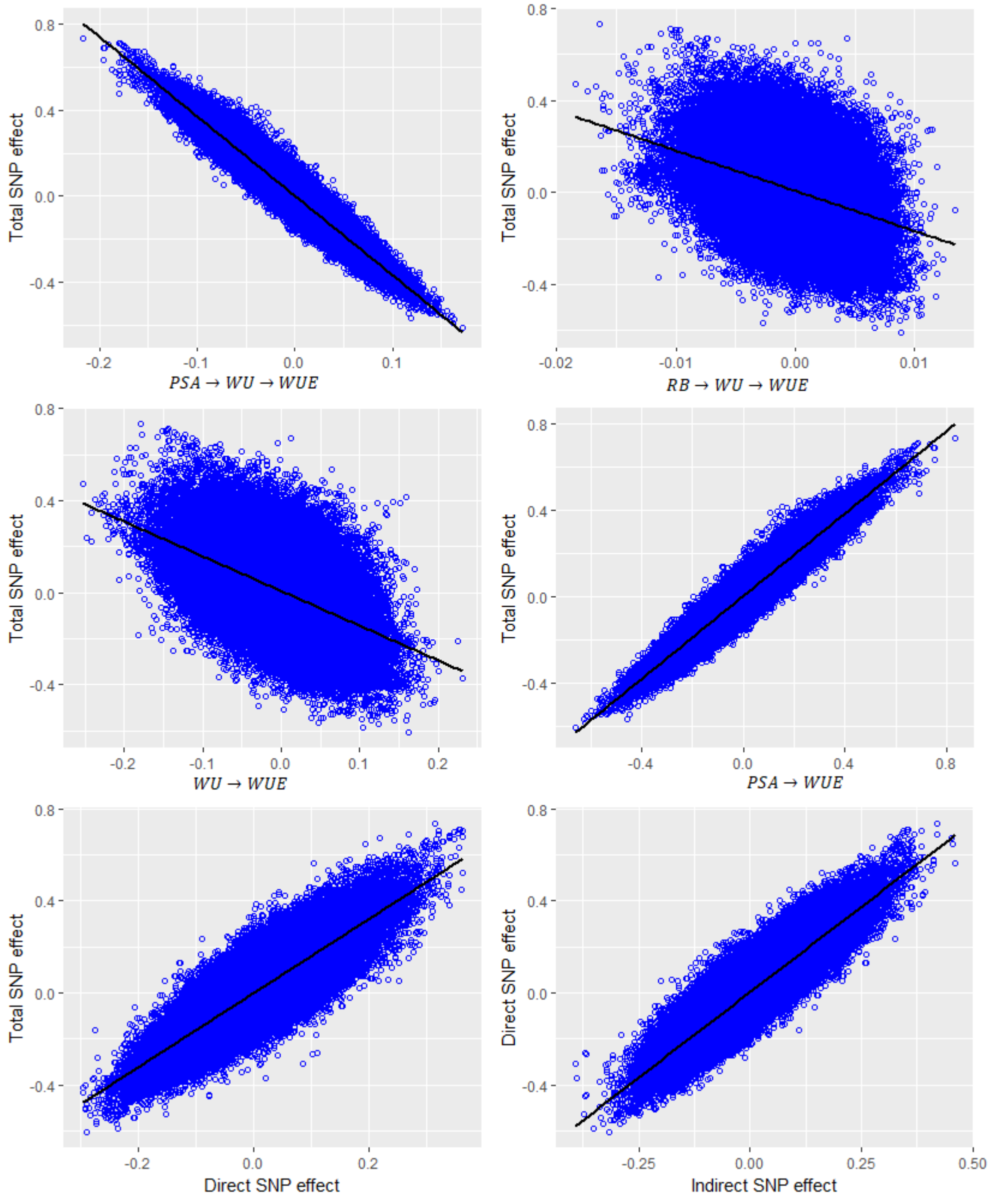


Figure S2: Scatter plots of decomposed SNP effects for water usage efficiency (WUE). Each point represents the estimated SNP effect for a SNP which affect WUE directly or indirectly. PSA: projected shoot area; RB: root biomass; WU: water use.

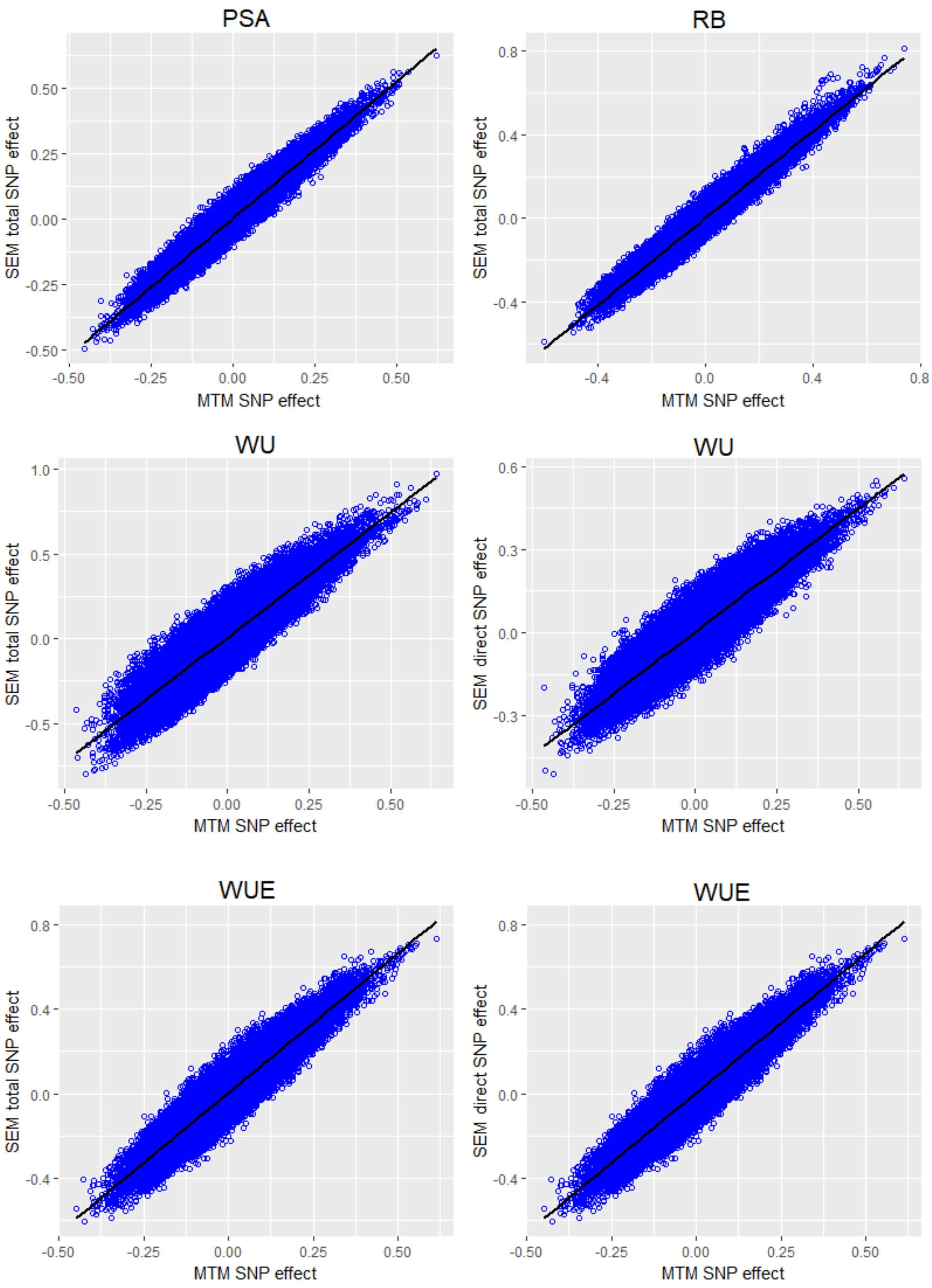


Figure S3: Scatter plots of estimated SNP effect between multi-trait model (MTM) and structural equation model GWAS. PSA: projected shoot area; RB: root biomass; WU: water use; WUE: water use efficiency.
